# Early AAV9-mediated *SLC9A6* gene replacement preserves cerebellar structure and motor function in the *shaker* rat model of Christianson syndrome

**DOI:** 10.64898/2026.11.04.764455

**Authors:** Kajal Dahiya, Kaanchana Sekaran, Lingye Hu, Andy Bloomfield, Evan Kapasakis, Sonja Henning-Smith, Lisa G Riley, Mactier Evans, Sarah Haslam, Joseph Katakowski, Daria Nesterovich Anderson, Wendy Gold, Collin J Anderson

## Abstract

Christianson syndrome (CS) is an X-linked neurodevelopmental and neurodegenerative disorder caused by *SLC9A6* loss of function, resulting in NHE6 deficiency and epilepsy, intellectual disability, cerebellar atrophy, and ataxia. Following on our prior gene replacement therapeutic development, we preclinically evaluated AAV9-mediated *SLC9A6* gene replacement in the context of promoter selection, dosing, and treatment timing. Intracerebroventricular delivery of neuronal-targeted gene replacement at 1 × 10¹¹ vector genomes (vg) produced substantial improvements in motor coordination and preserved cerebellar volume, quantified through structural MRI. Early postnatal treatment prevented the development of gait ataxia throughout the longitudinal assessment period, statistically indistinguishable from wildtype controls, with molecular analyses demonstrating restored NHE6 expression and preservation of cerebellar markers. Interestingly, later treatments at timepoints corresponding to early neurodegenerative stages yielded strong initial effects, but these effects were largely lost over time, with half-life modelling demonstrating the need for early treatment. This conflicted with more persistent motor benefit shown in prior work with an engineered capsid, suggesting the need to consider the interaction between treatment timing and capsid-specific tropism and/or persistence. Finally, we built models capable of accurately predicting motor coordination from several protein markers. These findings will impact therapeutic development for CS, with relevance across neurodevelopmental genetic disorders.

## Introduction

Christianson syndrome (CS) is a rare X-linked recessive disorder caused by mutations in the *SLC9A6* gene leading to sodium/hydrogen exchanger 6 (NHE6)^1,2^ loss of function. NHE6 is a membrane protein that regulates luminal pH levels within endosomal compartments, and NHE6 loss of function results in overacidification of the endolysosomal system, yielding a mixed neurodevelopmental and neurodegenerative pathology, with particular loss of Purkinje cells (PCs) of the cerebellum^3,4^. Resultingly, the core clinical manifestations of CS include severe intellectual disability, epilepsy, progressive gait ataxia, postnatal microcephaly, non-verbal status, hyperkinesia, and abnormal pain processing^1,2,5,6^. Clinical management for CS is currently limited to minimally effective symptomatic treatment with anti-seizure medications and supportive care.

In 2023, following decades of use as a naturally-occurring model of cerebellar degeneration and consequent ataxia and tremor^5–12^, the Wistar Furth *shaker* rat was identified as a naturally occurring rat model of Christianson syndrome, with a phenotype driven by a frameshifting, early-truncating mutation in *Slc9a6^13^*. Indeed, following identification *of Slc9a6* as a candidate gene, restoration of an average of ∼10% cerebellar NHE6 via a PC-targeted adeno-associated viral (AAV) vector improved both the motor phenotype and several forms of motor pathology^14^. Subsequent *shaker* rat-based analysis of the same Purkinje cell-targeted AAV and a more translationally relevant vector driving expression of human *SLC9A6* ubiquitously through the AAV9 serotype further supported AAV-mediated gene replacement as a therapeutic strategy for CS^15^.

AAVs, particularly AAV9, are now clinically established in the context of gene replacement for monogenic loss-of-function neurological disorders^16,17^. However, prior to any consideration of clinical gene therapy for CS, several key questions must be resolved, such as in the context of promoter selection, the temporal therapeutic window, treatment efficacy in the context of non-motor symptoms, and safety- and toxicity-related considerations. Thus, systematic evaluation of these parameters is necessary, and we sought to advance AAV9-mediated *SLC9A6* gene replacement closer towards clinical translation by systematically evaluating several of the above variables in the context of the *shaker* rat. We first compared several clinically relevant promoters in a dose-dependent and longitudinal manner in a cohort of 30 rats, evaluating degree of gene replacement and protection against gait ataxia and cerebellar atrophy as primary outcomes. Subsequently, we proceeded to evaluate the effects of varying the timepoint of administration of the identified lead vector, AAV9-SYN1-SLC9A6 in a second longitudinal study of a cohort of 37 rats.

Remarkably, we report a complete protection against motor phenotypic development through 25 weeks of age with early postnatal treatment with AAV9-SYN1-SLC9A6 at a dose of 1 × 10¹¹ vg/rat. Gene replacement rescued NHE6, preserved Purkinje cell health, and prevented cerebellar atrophy as quantified through structural MRI. Further, both cerebellar volume and several molecular markers were strongly associated with motor coordination. However, increases in ionized calcium-binding adapter molecule 1 (IBA1) expression in untreated Wistar Furth *shaker* rats was unresolved by gene replacement and inconsistent changes in glial fibrillary acidic protein (GFAP) highlight the importance of evaluating glial responses alongside therapeutic efficacy. Together, these findings provide critical insights into promoter selection, vector dosing and therapeutic timing for the development of gene replacement therapies for Christianson syndrome.

## Results

All experiments reported on occur in the context of Wistar Furth rats, both Wistar Furth *shaker* rats and wildtype Wistar Furth rats as controls. Male and female rats were utilised at random, with *shaker* rats referring to males that are hemizygous for the *shaker* mutation in *Slc9a6* and females that are homozygous for the *shaker* mutation; no heterozygous female rats are reported on. Thus, all *shaker* rats in this study feature total NHE6 loss of function and wild type rats have normal NHE6 function. Two separate longitudinal studies were carried out: 30 total rats were enrolled in a study to compare two promoters at two doses each, and second, 37 total rats were enrolled in a study to vary the timepoint of administration of the lead promoter from the first study at a high dose at 5 separate timepoints. Additionally, 8 additional rats were enrolled in MRI studies, leading to a total of 75 rats reported on.

### 3.1 Gene replacement reduces motor dysfunction in a dose-, promoter-, and timepoint-dependent manner

Gait ataxia was assessed longitudinally from 8 to 25 weeks of age in wildtype controls and *shaker* rats administered AAV9-SYN1-SLC9A6, AAV9-CAG-SLC9A6, or a AAV9-CAG-GFP control at doses of 1 × 10^11^ or 1 × 10^10^ vg, using established force plate actometry-based approaches^13,15,18,19^ (Fig. 1A). A linear mixed-effects model showed that ataxia trajectories across ages 8–25 weeks differed between groups (Joint Wald F(25, 78.1) = 29.98, p < .001) (Fig. 1B). Post-hoc testing demonstrated that, compared to untreated *shaker* rats, ataxia was reduced by AAV9-SYN1-SLC9A6 at 1 × 10^11^ vg (F(5,73.9)=23.08, p<.001), AAV9-SYN1-SLC9A6 at 1 × 10^10^ vg (F(5,69.6)=12.95, p<.001), and AAV9-CAG-SLC9A6 at 1 × 10^11^ vg (F(5,81.33)=2.94, p<.034). However, AAV9-CAG-SLC9A6 at 1 × 10^10^ vg did not significantly modify ataxia compared to untreated *shaker* rats. Finally, we evaluated the effects of promoter, dose, and their interaction among treated groups within the mixed model, finding a significant effect for each (SYN1 vs CAG: Wald F(5,84.613)=17.23, p<.001; 1E11 vg vs 1E10 vg: F(5,84.613)=3.51, p=.0063; dose v promoter interaction: F(5,84.613)=4.35, p=.0014).

**Figure 1:**
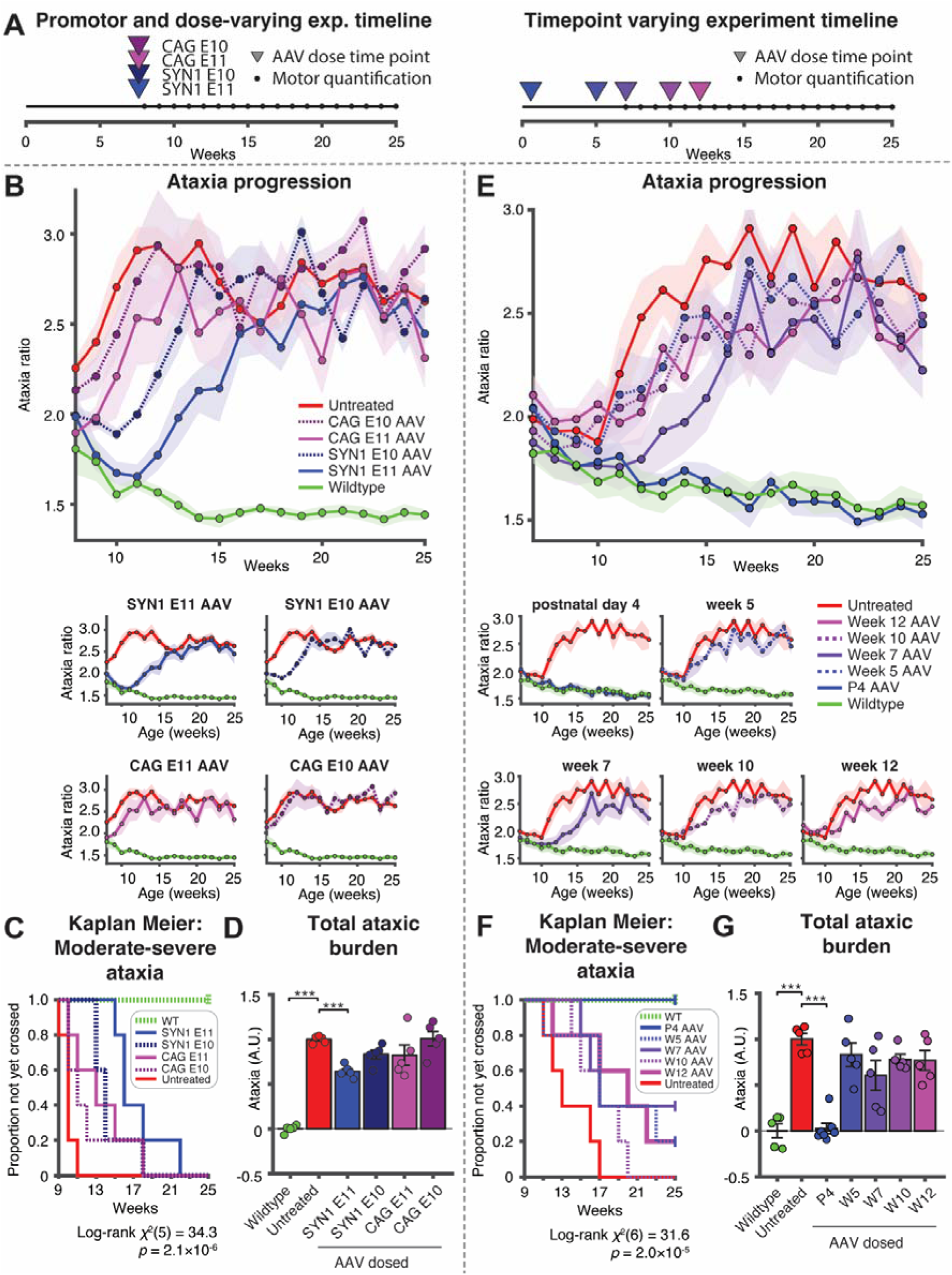
AAV9-mediated SLC9A6 gene replacement reduces gait ataxia. **A**. Experimental design for promoter/dose optimisation and treatment-timing studies. **B-D** show dose/promoter varying experimental data, while **E-G** show timepoint varying experimental data using AAV9-SYN1-SLC9A6 at 1 x 10^11^ vg. **B.** Longitudinal gait ataxia scores (8–25 weeks) in wildtype, untreated *shaker*, and *shaker* rats treated with AAV9-SYN1-SLC9A6 or AAV9-CAG-SLC9A6 at 1 x 10^10^ or 1 x 10^11^ vg, finding both a dose and promoter effect: SYN1 more effective than CAG in reducing ataxia, and 10^11^ vg more effective than 10^10^ vg. Lower panels show individual treatment groups compared to wildtype and untreated *shaker* rats. **C.** Kaplan–Meier analysis demonstrates a significant delay in reaching moderate-to-severe ataxia in all groups compared to untreated *shaker* rats except *shaker* rats treated with AAV9-CAG-SLC9A6 at 1 x 10^11^ vg. **D.** Total ataxia burden across promoter/dose groups, simplified from B shows a significant reduction in *shaker* rats treated with AAV9-SYN1-SLC9A6 compared to untreated *shaker* rats. **E.** Longitudinal gait ataxia scores (8–25 weeks) in wildtype, untreated *shaker*, and *shaker* rats treated with AAV9-SYN1-SLC9A6 1 x 10^11^ vg at time points ranging from P4 to 12 weeks of age, finding a correction of motor ataxia with P4 treatment to the point of insignificant difference from wildtype rats. **F.** Kaplan–Meier analysis demonstrates a significant delay in reaching moderate-to-severe ataxia in wildtype and P4-treated rats compared to untreated *shaker* rats and trends towards delay in all other groups. **G.** Total ataxia burden across promoter/dose groups, simplified from E shows a significant reduction in *shaker* rats treated at P4 compared to untreated *shaker* rats.

Next, we evaluated ataxic threshold crossings ranging from mild (ataxia ratio of 2.0) to very severe (ataxia ratio of 3.0) across groups. We found a significant group-wise difference at all thresholds (χ^2^(5)=34.25, p<.001 for moderate-severe threshold shown in Fig. 1C). Post-hoc testing for all other groups compared to untreated *shaker* rats in the moderate-severe threshold revealed a significant delay for each of wildtype (χ^2^(1)=9.53, p=.0019), AAV9-SYN1-SLC9A6 at 1E11 vg (χ^2^(1)=9.53, p=.0019), and AAV9-SYN1-SLC9A6 at 1E10 vg (χ^2^(1)=9.53, p=.0019). AAV9-CAG-SLC9A6 at 1E11 vg trended towards delay (χ^2^(1)=4.87, raw p=.027, p=.055 by Holm-Bonferroni correction), and CAG at 1E10 vg likewise trended towards delay (χ^2^(1)=2.83, p=.093).

As a simplified follow-up to the longitudinal analysis above, we finally analysed total ataxic burden from 8-25 weeks of age across all groups, finding a group-wise difference (F(5,24) = 35.70, p<.001) (Fig. 1D). All other groups were compared to untreated *shaker* rats, finding less ataxic burden in both wildtype rats (t=38.61, p<.001) and *shaker* rats treated with AAV9-SYN1-SLC9A6 at 1E11 vg (t=10.08, p<.001), with insignificant differences in other groups compared to untreated *shaker* rats.

Following dose- and promoter-varying studies, we assessed the relationship between treatment timing and gait ataxia with all treated rats receiving administration of AAV9-SYN1-SLC9A6 at 1 × 10^11^ vg at varying timepoints. As before, a linear mixed-effects model showed that ataxia trajectories differed between groups, although in this experiment we evaluated from 10-25 weeks of age given a slightly later onset in untreated *shaker* rats in this experimental cohort compared to the first longitudinal study (Joint Wald F(30, 87.80) = 14.26, p<.001) (Fig. 1E). In post-hoc testing comparing to untreated *shaker* rats, ataxia was lower in wildtype rats (F(5,90.41)=32.91, p<.001), as well as *shaker* rats treated at p4 (F(5,95.85)=34.64, p<.001), 7 weeks (F(5,88.65)=8.61, p<.001), 10 weeks (F(5,88.13)=3.56, p=.0056), and 12 weeks (F(5,86.91)=4.70, p<.001). *Shaker* rats treated at 5 weeks were insignificantly different from untreated *shaker* rats (F(5,90.44)=2.36, p=.046 raw, p=.092 corrected), and p4-treated *shaker* rats were insignificantly different from wildtype rats (F(5,93.31)=0.99, p=.430).

As above, we next evaluated ataxic threshold crossings ranging from mild (ataxia ratio of 2.0) to very severe (ataxia ratio of 3.0) across groups. We found a significant group-wise difference at all thresholds (χ^2^(5)=31.56, p<.001 for moderate-severe threshold shown in Fig. 1F). Post-hoc testing for all other groups compared to untreated *shaker* rats in the moderate-severe threshold revealed a significant delay for each of wildtype (χ^2^(1)=9.70, p=.002) and P4 treatment (χ^2^(1)=13.20, p<.001), while other treated groups trended towards delay (2.86≤χ^2^(1)≤4.32, 0.038≤raw p≤0.092, all corrected p>0.05).

Finally, as a simplified follow-up to the longitudinal analysis above, we once again analysed total ataxic burden from 10-25 weeks of age across all groups, finding a group-wise difference (F(6,30) = 20.19, p<.001) (Fig. 1G). All other groups were compared to untreated *shaker* rats, finding less ataxic burden in both wildtype rats (t=10.90, p<.001) and *shaker* rats treated with AAV9-SYN1-SLC9A6 at 1E11 vg (t=12.32, p<.001), with insignificant differences in other groups compared to untreated *shaker* rats.

### 3.2. Gene replacement restores NHE6 expression with strong timepoint dependence

Following the longitudinal motor experiments, we proceeded to characterise the molecular effects of the SYN1-based AAV strategy. Thus, we started by quantifying cerebellar NHE6 expression via western blotting in cerebellums collected from ∼30-week-old rats following administration of AAV9-SYN1-SLC9A6 1 × 10¹¹ vg at P4 or 5, 7, 10 and 12 weeks of age compared to age-matched wildtype and untreated *shaker* rats (Fig. 2). First, we found that NHE6 expression was significantly reduced in untreated *shaker* rats compared to wildtype rats (t=10.71, p<.001). Next, we compared across all 7 groups, finding that NHE6 expression differed significantly across groups (*F*(7, 36) = 14.93, *p* < .001, η² = 0.74). Further, *shaker* rats treated at P4 had greater NHE6 than both untreated *shaker* rats (t=8.19, p<.001) and wildtype rats (t=2.54, p=.016). Further, *shaker* rats treated at 5, 7, and 10 weeks of age expressed greater NHE6 than untreated *shaker* rats (5W: t=5.85, p=.0021; 7W: t=4.76, p=.0044, 10W: t=6.24, p=.0016), but a significant difference was not found between *shaker* rats treated at 12 weeks of age and untreated *shaker* rats.

**Figure 2:**
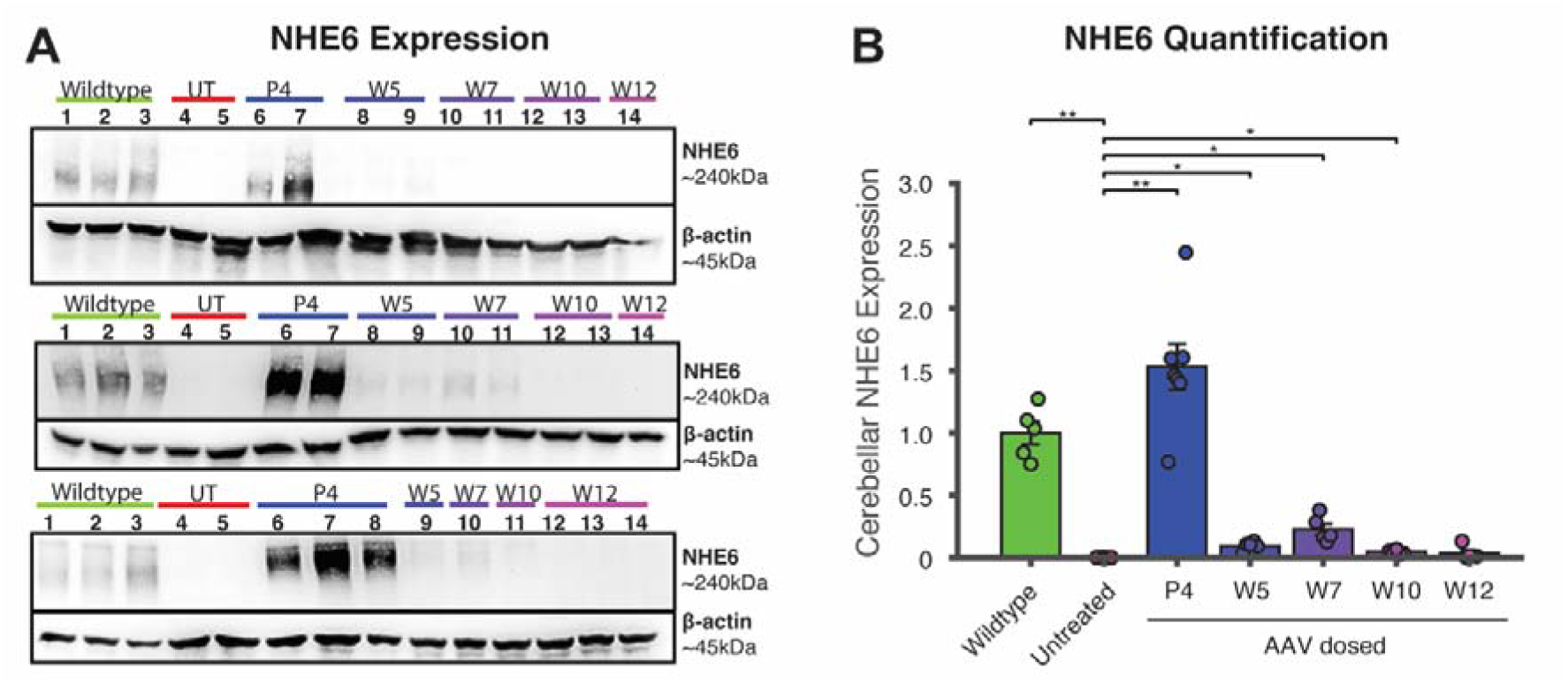
AAV9-mediated neuronally targeted *SLC9A6* gene replacement restores cerebellar NHE6 expression, particularly when administered early. **A.** Representative western blots showing cerebellar NHE6 expression across three blots in wildtype (WT), untreated *shaker* (UT) and AAV9-SYN1-SLC9A6-treated *shaker* rats administered at postnatal day 4 (P4) or weeks 5, 7, 10 and 12. Each blot included three WT and two UT reference lanes, with treatment samples distributed across blots (n = 5 biologically independent animals per group; 7 groups, n = 35). **B.** NHE6 expression was normalised to β-actin and expressed relative to the mean WT expression. Repeated WT measurements were averaged before normalisation. P4-treated *shaker* rats exhibited greater NHE6 expression than both wildtype rats and untreated *shaker* rats, while treatment of *shaker* rats at each of W5, W7, and W10 led to small but significant increases in NHE6 compared to untreated *shaker* rats and treatment at W12 did not significantly modify cerebellar NHE6 expression

### 3.3. Gene replacement corrects several Purkinje cell molecular deficits

We quantified PCP2, CALB1, and RGS8 expression via western blotting in cerebellums collected from ∼30 week-old rats administered AAV9-SYN1-SLC9A6 1 × 10¹¹ vg at P4 or 5, 7, 10 and 12 weeks of age (Fig. 3A-B) compared to age-matched wildtype and untreated *shaker* rats. PCP2 (Fig. 3A-B) (t=12.46, p<.001), CALB1 (Fig. 3C-D) (t=5.65, p=.0024), and RGS8 (Fig. 3E-F) (t=5.64, p=.0022) expression was significantly reduced in *shaker* rats compared to wildtype rats. Next, we next compared across all 7 groups, finding significant group-based differences for each (PCP2: F(6, 30) = 31.11, p < .001, η² = 0.86; CALB1: F(6, 30) = 22.38, p < .001, η² = 0.82; RGS8: F(6, 30) = 14.63, p < .001, η² = 0.75). Further, compared to untreated *shaker* rats, P4-treated *shaker* rats showed greater expression of PCP2 (t=6.23, p<.001), CALB1 (t=5.86, p<.001), and RGS8 (t=8.16, p<.001). Finally, *shaker* rats treated at 12 weeks were found to have greater CALB1 (t=3.59, p=.0052) and RGS8 (t=2.97, p=.0091) expression than untreated *shaker* rats. Notably, CALB1 was initially found to be greater in wildtype rats than P4-treated *shaker* rats, as was PCP2 in week 7-treated *shaker* rats compared to untreated *shaker* rats, but statistical significance was lost in each with the application of a Holm-Bonferroni correction.

**Figure 3:**
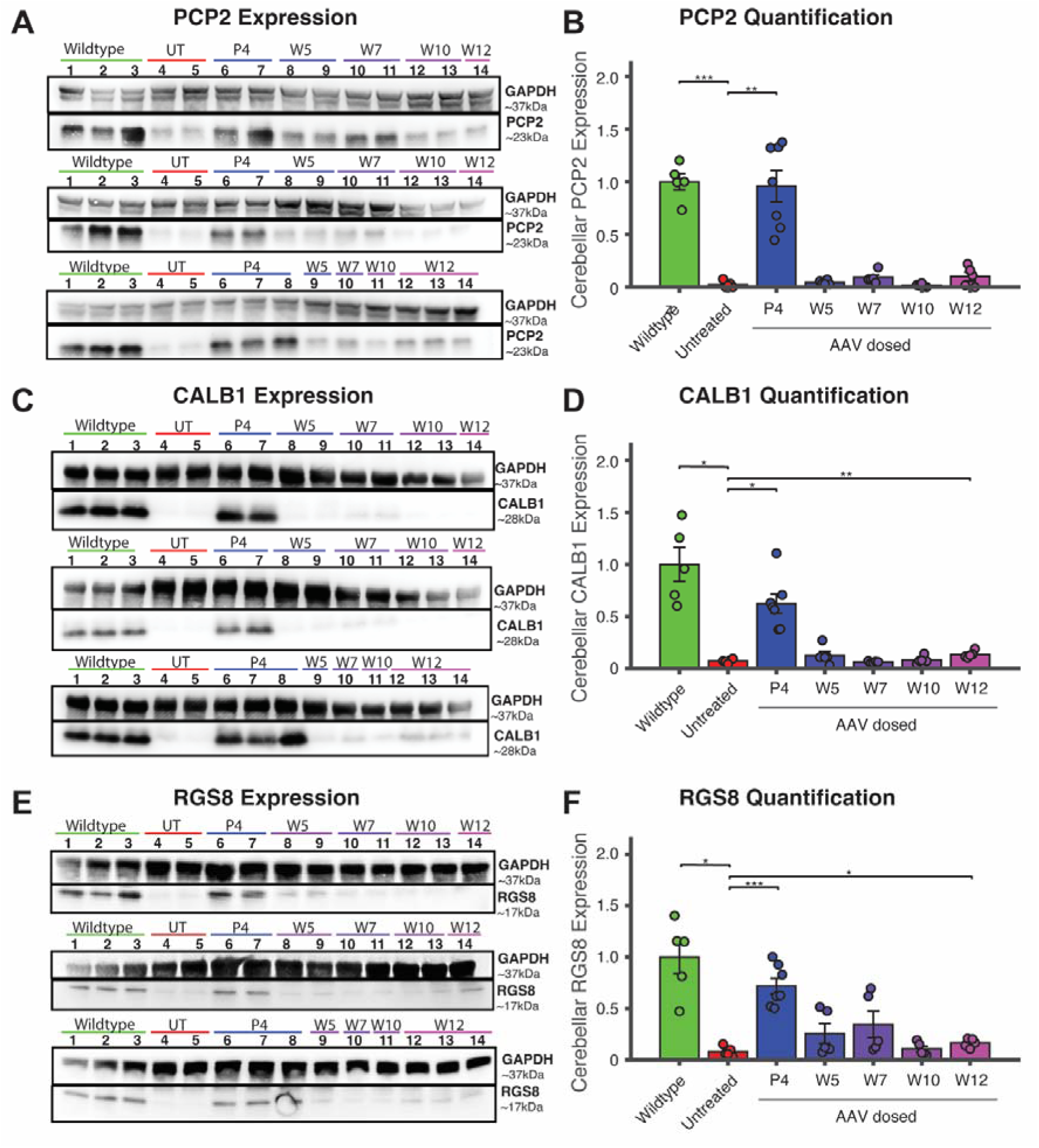
Early postnatal AAV9-mediated *SLC9A6* gene replacement restores Purkinje cell-associated protein expression. Western blots show cerebellar PCP2 (**A**), CALB1 (**C**), and RGS8 (**E**) expression, respectively, in wildtype (WT), untreated *shaker* (UT) and AAV9-SYN1-SLC9A6-treated *shaker* rats (P4, W5, W7, W10 and W12). **B. D. F.** Corresponding protein quantification normalised to β-actin and expressed relative to mean WT expression. Most notably, P4-treated *shaker* rats were found to have significant increases in each protein compared to untreated *shaker* rats, all insignificantly different from WT.

### 3.4 Gene replacement reduces cerebellar atrophy

Hemicerebellar volume was quantified via structural 7T MRI post-mortem in rats following both longitudinal studies (Fig. 4A-B). Across both cohorts, hemicerebellar volume was strongly correlated with late-stage ataxia, averaged across the final 5 weeks of the longitudinal motor study and predicted via a linear model accounting for both sex and, given the differential timepoint of tissue collection across experiments, the experimental enrolment (Fig. 4C) (partial r=-.73, p<.001). Hemicerebellar volumes were first compared between wildtype rats and untreated *shaker* rats following both longitudinal experiments, with reduced hemicerebellar volume in untreated *shaker* rats compared to wildtype rats in both the dose/promoter study (t=14.60, p<.001) and the timepoint varying study (t=5.88, p<.001) (Fig. 4D). In the dose/promoter varying study, hemicerebellar volume differed significantly across groups (F(5, 23) = 22.89, p < .001, η² = 0.83). In the dose and promoter variation experiment, we quantified group differences and found a significant group-based difference (F(5, 23) = 22.89, p < .001, η² = 0.83). Post-hoc tests showed that *shaker* rats treated with AAV9-SYN1-SLC9A6 at 1E11 vg had significantly larger hemicerebellums than untreated *shaker* rats (t=3.60, p=.0053), while other comparisons to untreated did not yield significant differences. In the timepoint variation experiment, we quantified group differences and found a significant group-based difference (F(5, 30) = 11.92, p < .001, η² = 0.67). Post-hoc tests showed that P4-treated *shaker* rats had significantly larger hemicerebellums than untreated *shaker* rats (t=4.64, p<.001), while other comparisons to untreated (as well as P4-treated *shaker* rats vs WT) did not yield significant differences.

**Figure 4:**
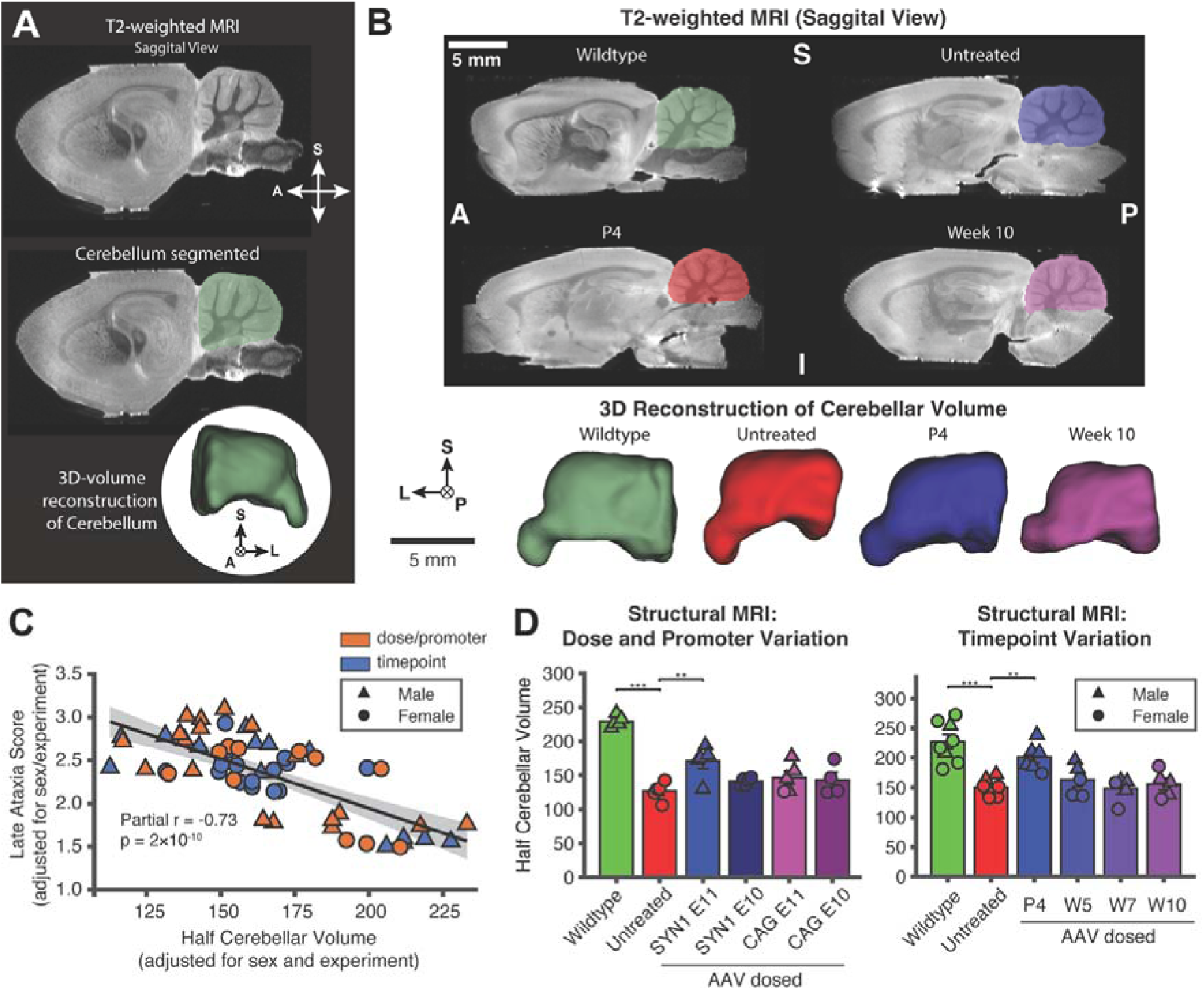
Neuronally-targeted *SLC9A6* gene replacement attenuates cerebellar atrophy in *shaker* rats. **A.** Representative MRI scans illustrating hemicerebellar segmentation. **B.** Representative three-dimensional reconstructions of hemicerebellar volumes across groups in the timepoint variation experiment. **D.** L: Hemicerebellar volume (mm³) in wildtype (WT), untreated shaker (UT) and SYN1- or CAG-treated shaker rats receiving 1 × 10¹ or 1 × 10¹¹ vg. R: Hemicerebellar volume following AAV9-SYN1-SLC9A6 (1 × 10¹¹ vg) administration at P4, W5, W7, W10.

### 3.5. Gene replacement does not drive clear autophagic or glial effects

Next, we quantified cerebellar p62, GFAP, and IBA1 expression via western blotting in cerebellums collected from ∼30 week-old rats administered AAV9-SYN1-SLC9A6 1 × 10¹¹ vg at P4 or 5, 7, 10 and 12 weeks of age (Fig. 5) compared to age-matched wildtype and untreated *shaker* rats. First, we compared expression of each between wildtype rats and untreated *shaker* rats, finding that untreated *shaker* rats exhibited greater p62 (t=2.33, p=.039) and IBA1 (t=3.84, p=.0058), while no significant difference was found in GFAP. Given a lack of difference in wildtype vs untreated *shaker* rats for GFAP, we proceeded with group-wise difference for only p62 and IBA1. A strong trend was found for a group-wise difference for p62, but it was not statistically significant (F(6, 30) = 2.18, p = .073, η² = 0.3); thus, post-hoc testing was not carried out for p62. Likewise, a strong trend was found for a group-wise difference for IBA1, but it was not statistically significant (F(6, 30) = 2.29, p = .062, η² = 0.31); as with p62, post-hoc testing was not carried out for IBA1. However, a trend was noted that treated groups demonstrated numerically more IBA1 expression than untreated *shaker* rats.

**Figure 5:**
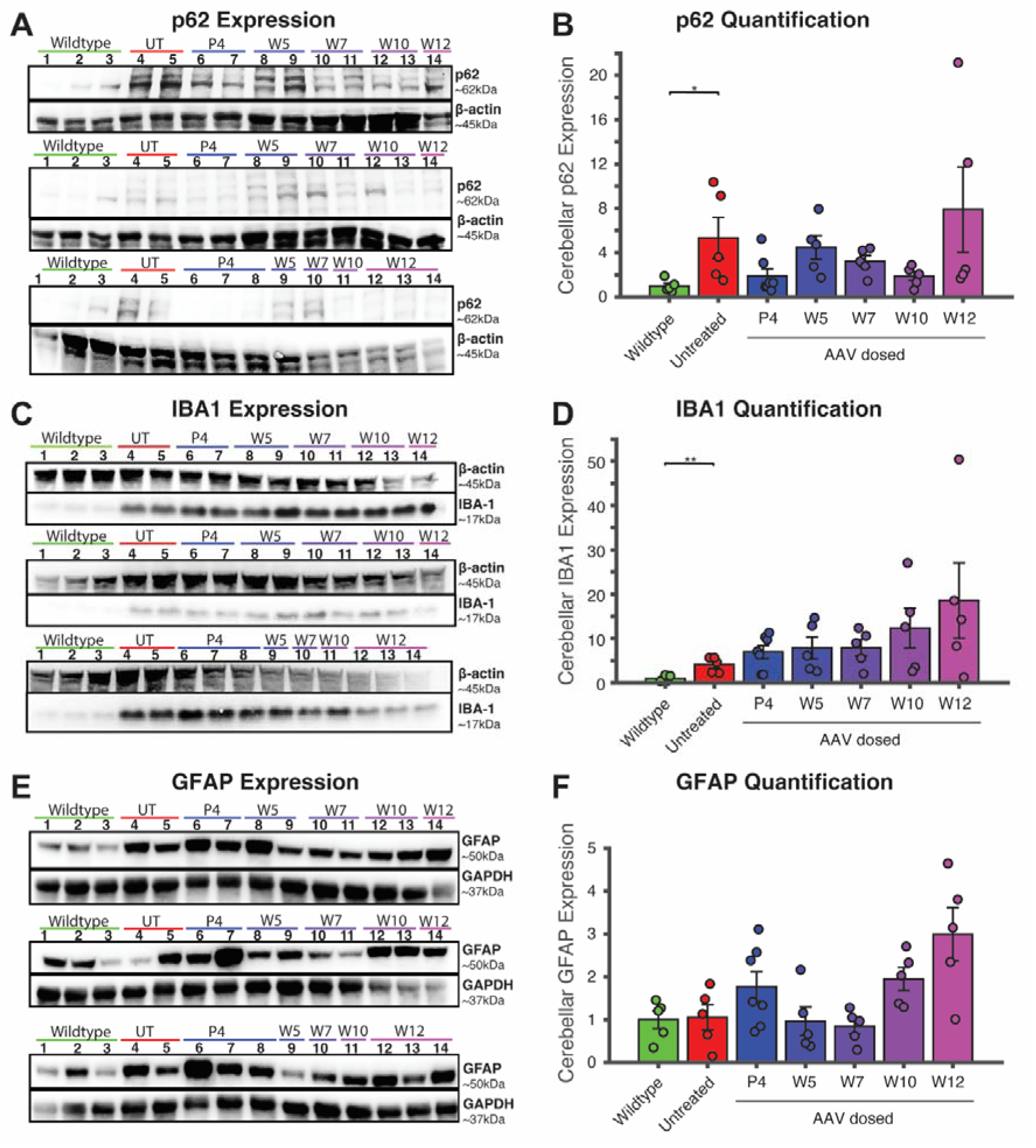
Autophagic and microglial reactive changes are found in untreated *shaker* rats compared to wildtype controls, but a significant treatment effect was not found. **A. C. E**. Representative western blots showing cerebellar p62, IBA1, and GFAP expression, respectively, in wildtype (WT), untreated *shaker* (UT) and AAV9-SYN1-SLC9A6-treated *shaker* rats (P4, W5, W7, W10 and W12). **B. D. F.** Corresponding protein quantification normalised to β-actin or GAPDH and expressed relative to mean WT expression. Early postnatal treatment was associated with reduced p62 expression compared to untreated *shaker* rats (∼80% reduction towards wildtype rats, p=0.070), although statistical significance was not found. This may indicate partial restoration of autophagic degradation with early gene replacement, although direct measures of autophagic flux would be needed for confirmation.

### 3.6 Early treatment is crucial to optimise AAV9-based efficacy due to short half-lives of treatment timing in motor, structural, and molecular outcomes

Having quantified motor, structural MRI, and several molecular responses to differentially timed treatment with AAV9-SYN1-SLC9A6, we next characterised the relationship between dose timing and each outcome using the 5 dose time points from P4 to W12. We z-scored ataxia outcomes across all animals, and we calculated linear slopes against dosing age (adjusted for sex) to quantify how each 1-week delay in AAV dosing time affected each marker’s quantification. With every week in delay in AAV administration, ataxia motor scores increased by 0.15 SD/week (p < 0.001), while NHE6, PCP2, CALB1, RGS8, and MRI volume also decreased (range: -0.204 – -0.113SD/week, p < 0.001) (Fig. 6A). For motor, NHE6, PCP2, CALB1, and RGS8 metrics, exponential fits better captured the decay from wildtype to untreated marker levels (Fig 6B). For MRI volume decay, the exponential and linear model were comparable. Generally, we observed that dosing at weeks 5, 7, 10, and 12 already showed low endpoint rescue percentage, indicating that early AAV administration is critical (Fig 6B). In contrast, markers p62, IBA1, and GFAP trended towards increase post-AAV administration, so an exponential decay model was not appropriate. Instead, we quantified how each marker increased as a multiplicative factor over wildtype levels (Fig 6C). For p62, untreated *shaker* rats had 4.12 times more p62 than wildtype rats, but the increasing trend with dosing age, assessed on a log_2_ scale, was not statistically significant. Notably, IBA1 and GFAP levels in later-dosed AAV groups trended above both untreated and wildtype levels, raising the possibility of a vector-associated glial response, although this slope likewise did not reach significance for either marker.

**Figure 6:**
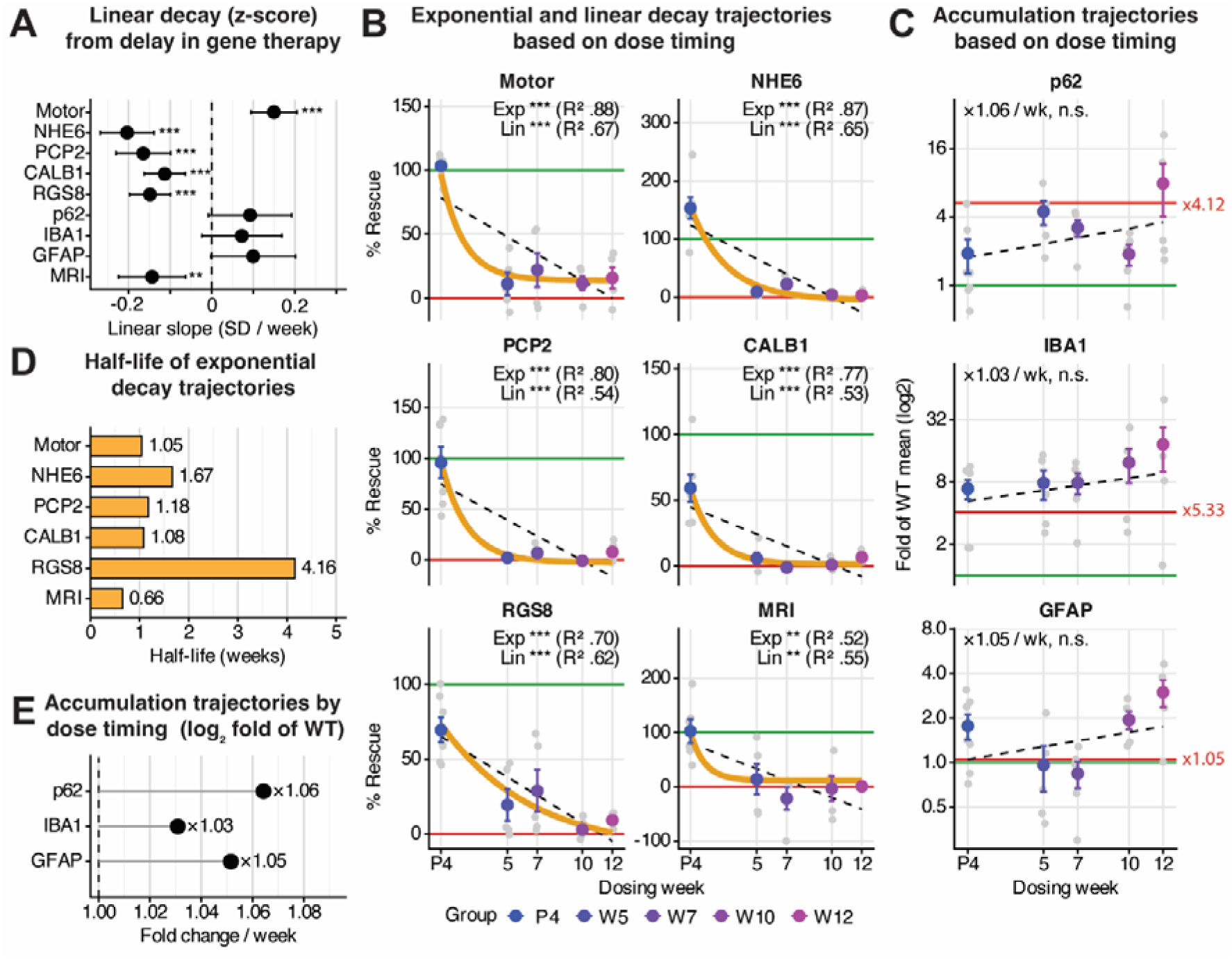
Delayed administration of AAV9-SYN1-SLC9A6 results in a rapid loss of late-stage motor, molecular, and structural protection. **A.** With each additional week of delay in AAV administration, late-stage NHE6, PCP2, CALB1, and RGS8 significantly declined (range: -0.204 – -0.113 SD/week, p<0.001), MRI volume significantly declined (-0.143 SD/week, p<0.01), and motor scores significantly increased, indicating worsened ataxia (0.15 SD/week, p<0.001). **B.** Changes in Motor, NHE6, PCP2, CALB1, and RGS8 with respect to dose timing are better captured by an exponential decay model than a linear model. Expression values are normalised from untreated at 0% (red horizontal line) to wildtype at 100% (green horizontal line). **C.** Changes in p62, IBA1, and GFAP were expressed as fold change relative to the WT mean (1.0, green horizontal line), and we determined weekly change by fitting a regression on a log2 scale with sex as a covariate. While we previously found that p62 (Fig 5B) and IBA1 (Fig 5D) were significantly increased in untreated versus wildtype cohorts, the increasing fold relationships of these markers with later AAV dose timing were not significant. Interestingly, IBA1 and GFAP had regression lines that exceeded untreated *shaker* levels, while the regression of p62 lay between wildtype and untreated levels. This suggests that GFAP and IBA1 may be increased by viral administration, while p62 changes may represent a treatment effect, with studies underpowered to detect significance. **D.** We extracted half-life values from exponential fits in B to quantify the delay in AAV administration resulting in a halving of metrics. Half-lives were less than 1 week for MRI volume (0.66 weeks, p < 0.01) and less than 2 weeks for Motor, NHE6, PCP2, and CALB1 (range: 0.66–1.67 weeks, p<0.001). RGS8 had the longest half-life of 4.16 weeks (p<0.001). **E.** A delay in AAV administration by n weeks increases marker levels by a multiple of FCⁿ, where FC is the fold change per week. Thus, a delay of 5 weeks in AAV administration for p62, represents an increase by 1.06^5^ (33.8%) in p62 relative to earlier treatment. Fold change relationships were largely trending but not significant (p62: p=0.130, IBA1: p=0.450, GFAP: p=0.15).

For decay trajectories in Fig. 6B, we quantified the theoretical half-life of each marker, representing a 50% loss in that marker’s quantification for every half-life delay in AAV administration (Fig 6D). The half-life values across markers are strikingly short: MRI volume has the shortest half-life of 0.66 weeks, while motor, NHE6, PCP2, and CALB1 showed a half-life of 1-2 weeks (range: 1.05–1.67 weeks); RGS8 had the longest half-life of 4.16 weeks. Thus, in the context of a rat, delaying AAV by a week has dramatic consequences on rescue of healthy phenotype. Accumulation trajectories showed only a modest, non-significant fold increase with later dosing (3–6% per week of delay) (Fig. 6E). Thus, whilst the comparison of all AAV-treated *shaker* rats to untreated *shaker* rats showed that AAV-treated rats had significantly higher IBA1 (Welch’s t=3.05, p=.0024) and a trend towards increased GFAP (Welch’s t=1.79, p=.054), whether such a potential elevation depends on dosing age could not be established.

### 3.7 Terminal molecular markers and sex explain nearly all variability in late-stage motor outcomes

In the context of AAV9-SYN1-SLC9A6 administration, we compared all molecular markers to late-stage ataxia, averaged across the final 5 weeks of the longitudinal experiment. We found that PCP2, NHE6, CALB1, and RGS8 had significant, inverse correlations with motor scores in the last 5 weeks of the experiment across all groups (Fig 7A; r=-0.86, r=-0.84, r=-0.79, and r=-.072, respectively; p<0.001). However, PCP2, CALB1, RGS8, and NHE6 were also all significantly correlated with each other (r = 0.65–0.84; p<0.001)—though this was anticipated given that they are direct markers of Purkinje health^20^.

**Figure 7:**
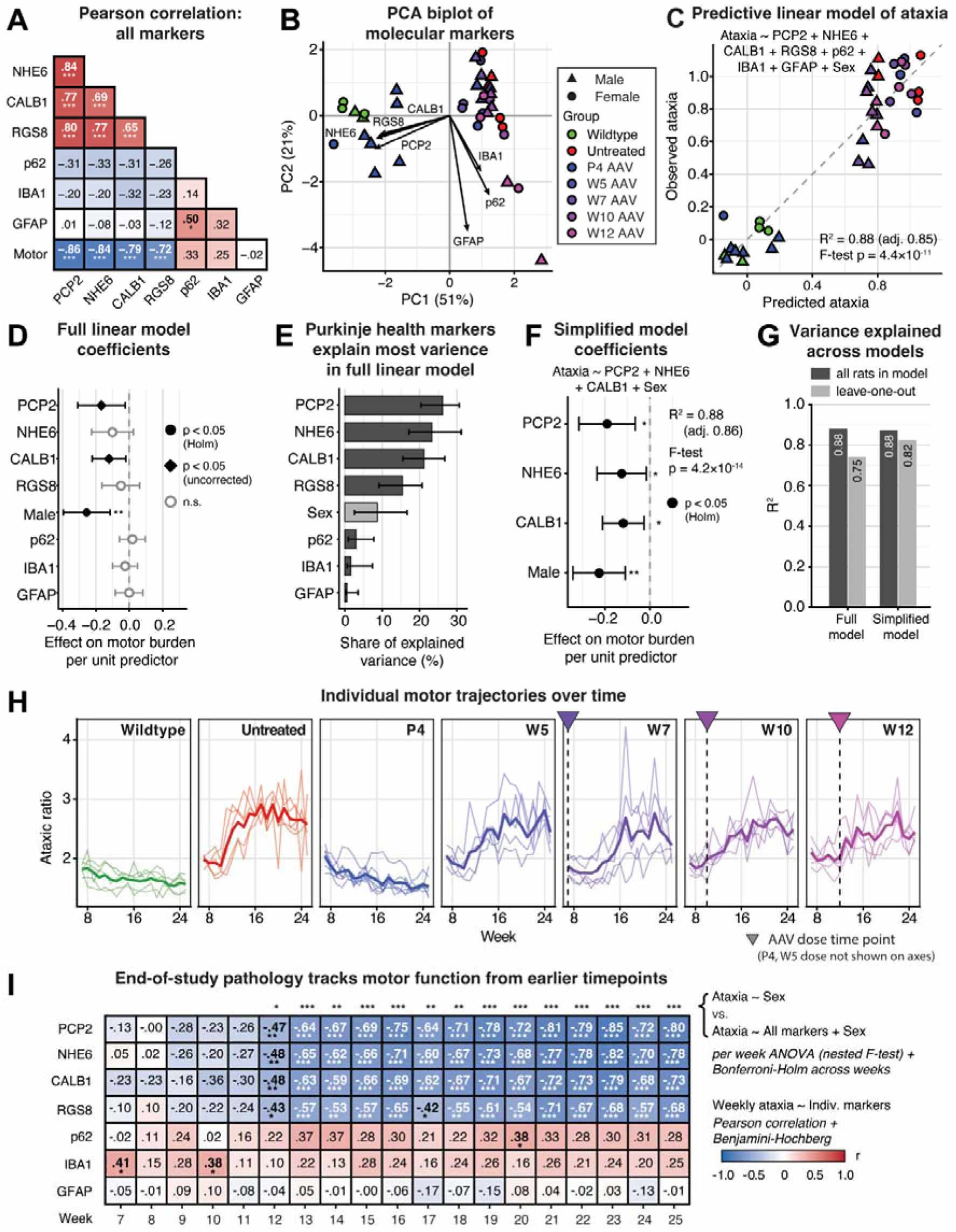
Robust relationship of ataxia with end-of-study molecular markers and sex as variables. **A.** Repeated Pearson correlations with Holm-Bonferroni post-hoc across all 8 markers demonstrate that NHE6, PCP2, CALB1, and RGS8 are positively correlated with each other (p< 0.001) and negatively correlated with end-of-study ataxia scores from weeks 21-25 (p<0.001). **B.** A principal component analysis biplot reveals NHE6, PCP2, CALB1, and RGS8 carry similar information in PCA1, which explains 51% of the variance in end-of-study ataxia scores. **C.** The full model containing all molecular markers and sex as variables can effectively predict end-of-study ataxia burden (R^2^ = 0.88, p = 4.4 x 10^-11^). **D.** In the full multivariate linear model in C, only sex was a significant predictor of ataxic burden (β=-0.23, p<0.01), while PCP2 and CALB1 coefficients were only significant before Holm-Bonferroni post-hoc correction. **E.** We ranked the variance each marker explained in the full model to build a simplified model with the top 3 markers (PCP2, NHE6, and CALB1) plus sex. **F.** The simplified model using only PCP2, NHE6, CALB1, and sex effectively predicted end-of-study ataxia burden (R^2^=0.88, p<.001). **G.** The performance of the simplified model is as good as the full model (R^2^=0.88 in both cases), but the simplified model performs better during leave-one-out-cross-validation (R^2^=0.82 versus R^2^=0.75). **H.** Visualised mean and individual trajectories of ataxic burden over time demonstrate increasing trajectories for untreated, W5, W7, W10, W12 cohorts. **I.** The heatmap shows repeated Pearson correlations for each of the 7 end-of-study molecular markers and weekly ataxic burden. After Benjamini-Hochberg post-hoc correction to ensure false discovery rate at q = 0.05, NHE6, PCP2, CALB1, and RGS8 emerged as significant predictors from week 12 onwards. Additionally, we found that also starting at week 12, the full multivariate model using sex and all molecular markers was significantly better at predicting weekly ataxic burden than a baseline model with sex alone, indicating week 12 is a critical timepoint where cohorts begin to diverge.

We generated a principal component analysis biplot, which demonstrated that these Purkinje and cell markers carried similar information along the first principal component (51% of the motor variance), while the inflammatory markers GFAP, IBA1, and p62 substantially contributed to the second principal component (21% of the variance) (Fig. 7B). To disentangle the individual effects of the markers, as well as considering sex as a variable, we built a multivariate linear regression using all markers and sex to predict motor outcomes. The model was highly predictive of motor scores, with an R^2^ of 0.88 (F-test, p = 4.4 x 10^-11^) (Fig. 7C) and performed better than any individual variable alone (R^2^ of 0.74 for Ataxia ∼ PCP2; Pearson). However, with seven variables in the model and despite collinearity between variables at acceptable levels, the model could not confidently differentiate each marker’s specific effect on motor burden. While PCP2, CALB1, and sex were initially significant variables in the model (p<0.05; β = -0.17, β = -0.12, and β = -0.26, respectively), only sex passed post-hoc correction (p= 0.000865 (adj. p=0.00692) (Fig 7D). Male rats had significantly lower ataxia scores than female rats, likely due to a heavier weight of males facilitating a greater signal-to-noise ratio, rather than biology difference in phenotype. We used relative importance analysis to isolate the top 3 markers (Fig 7E) and ran a simplified linear model using only PCP2, NHE6, CALB1, and sex as predictor variables. The simplified model performed nearly as well to the full model (R^2^ = 0.88) and all predictors passed post-hoc correction (p<0.05: PCP2 β = -0.19, CALB1 β = -0.12, NHE6 β = -0.12, and p<0.01: sex β = -0.23) (Fig. 7F). Both the full and simplified model passed leave-one-out-validation (p=1.32 x 10^-12^ and 1.75 x 10^-14^, respectively), but the larger drop in R^2^ for the full model in the leave-one-out approach (Fig. 7G) suggests greater dependence on individual animals in the full model, and that additional markers did not add predictive value beyond sex, PCP2, CALB1, and NHE6.

We next evaluated how molecular quantifications from tissue taken at the end of the experiment track ataxic burden at each of the weekly time points. Individual and cohort average ataxia trajectories sampled weekly are re-visualised in Fig. 7H. Finally, we conducted repeated Pearson correlations (Benjamini-Hochberg post-hoc) between weekly motor scores and each of the 7 molecular markers (Fig. 7I). Starting from week 12, each cerebellar marker (PCP2, CALB1, RGS8) and NHE6 were significantly negatively correlated with weekly ataxia measurement at all timepoints, and motor outcomes became more significantly correlated with each above-mentioned endpoint marker over the duration of the study (-.883<r<-.841, p<.001 for all). The joint contribution of all molecular markers to weekly ataxia was assessed by comparing a full model (markers + sex) against a reduced model (sex only), and after Bonferroni-Holm post-hoc correction, we found the full model became significant at the 12-week time point, indicating ataxia significantly diverged between groups at this point.

## Discussion

A vast majority of CS cases are caused by loss-of-function mutations in *SLC9A6* and consequent NHE6 loss of function, implicating gene replacement as a therapeutic strategy. Thus, we longitudinally evaluated promoter selection, vector dose, and treatment timing for AAV9-mediated *SLC9A6* replacement in the *shaker* rat model of CS. We found that early postnatal administration provided the greatest therapeutic benefit, with high-dose, neuronally targeted expression preventing the development of gait ataxia through 25 weeks of age, substantially reducing cerebellar atrophy, and markedly improving molecular markers of cerebellar pathology. Further, our findings identify treatment timing as a critical determinant of AAV9-based therapeutic efficacy, with quick loss of efficacy with delayed treatment dosing. Finally, we found that nearly all variability in preclinical CS ataxia can be explained through just 4 molecular markers.

SYN1 driven *SLC9A6* expression produced the strongest therapeutic benefit across the motor phenotype i.e. gait ataxia and structural endpoints among the promoter-dose combinations tested, although it must be stressed that SYN1-vs CAG-based vectors differed on more than just promoter. Indeed, due to packaging limitations and the greater size of the CAG promoter compared to the SYN1 promoter, the CAG-based vector did not include a woodchuck hepatitis virus posttranscriptional regulatory element, and thus, had its inclusion been possible, it is possible that CAG-based benefit may have matched or exceeded that of SYN1. Notably, in this comparative study, the treatment was initiated at ∼7.5 weeks of age, instead of ∼5-week ICV injections used in previous studies testing gene replacement therapy in *shaker* rats^13,15^. Despite the delayed treatment, AAV9-SYN1-SLC9A6 at 1 x 10^11^ vg delayed the onset of ataxia and substantially improved total ataxic burden.

The effect of treatment timing on motor phenotype following AAV9-SYN1-SLC9A6 1 10 vg is markedly non-liner. Rather than a gradual decline in efficacy with increasing age of administration, treatment at post-natal day 4 resulted in a disproportionately strong therapeutic response that resembled and was modelled well by an exponential decay function, mirroring prior work^21^. This suggests that the earliest postnatal period represents a distinct therapeutic window for AAV9-mediated correction, rather than simply the earliest point along a continuous age-dependent gradient. Several potentially overlapping mechanisms may contribute to this effect. First, AAV9 exhibits age-dependent changes in tropism, with neonatal administration favouring neuronal transduction more strongly than delivery later in development, when nervous system transduction shifts increasingly towards glial cells^22,23^. Early intracerebroventricular delivery may also enable broader vector distribution throughout the developing cerebellum before maturation of ventricular and parenchymal architecture limits access to target regions^23–25^. The developmental state of the cerebellum may provide another explanation. Cerebellar granule cell precursors undergo extensive postnatal proliferation^26^, with granule cell production continuing through the first two weeks in rodents^27^. At postnatal day 4, before substantial expansion of this population, Purkinje cells would therefore represent a relatively greater proportion of neuronal targets accessible to the vector. Given the prominent vulnerability of Purkinje cells in *SLC9A6* deficiency, this developmental context may increase the likelihood of transducing neurons that are particularly important for therapeutic benefit. Consistent with this possibility, neonatal AAV9 administration has been reported to achieve robust Purkinje cell transduction^23,28^.

Restoration of NHE6 was accompanied by preservation of multiple Purkinje cell-associated proteins and of cerebellar volume, supporting a broader protective effect on the cerebellar phenotype. CALB1, PCP2 and RGS8 are all strongly expressed in Purkinje cells and have previously been used as molecular markers of Purkinje-cell integrity in degenerative cerebellar disease. ^29,30,31,32,33^. As CALB1, PCP2 and RGS8 are expressed throughout substantial regions of the Purkinje-cell, including dendrites, reduced abundance may reflect a combination of neuronal loss, dendritic atrophy and altered protein expression within surviving cells, and thus should not be interpreted in the context of Purkinje cell quantities. Notably, preservation of cerebellar volume on MRI strengthens the interpretation that NHE6 replacement protects cerebellar tissue more broadly, although volumetric preservation likewise cannot distinguish neuronal survival from preservation of dendritic, glial or other tissue compartments.

Cerebellar volume loss and progressive Purkinje cell degeneration have previously been demonstrated in *Slc9a6*-deficient models^13,34^, and this work extends these findings. Notably, however, in the initial promoter and dose optimisation experiment, high-dose SYN1-driven gene replacement was associated with greater cerebellar volume relative to untreated *shaker* rats despite a loss of motor protection by late in the longitudinal study. Sex-based differences were not found in hemicerebellar volume in the timepoint varying study. However, it should not be ruled out a sex-based difference is still a plausible interpretation for the greater cerebellar volume in high-dose SYN1 gene replacement compared to untreated *shaker* rats, especially given the loss of motor protection over time.

The *shaker* phenotype was associated with a 400%+ increase in p62 accumulation, consistent with previous studies demonstrating impaired lysosomal function, reduced autophagic flux, and p62 accumulation in NHE6-deficient neurons^35,36^. AAV9-mediated *SLC9A6* replacement did not yield a statistically significant decrease in p62; however, P4-treated rats averaged a 79% reduction from untreated *shaker* p62 levels towards wildtype levels (p=.070, uncorrected) despite NHE6 expression. Thus, studies may simply have been underpowered to detect AAV-mediated p62 correction. However, p62 abundance alone would be unable to establish restoration of autophagic flux, and direct functional assessment would be required to determine normalisation of autophagy. CNS-directed AAV administration can elicit immune responses involving astrocytes and microglia, depending on vector characteristics and transgene immunogenicity^37,38^. Consistent with a potential treatment-associated response, IBA1 abundance generally trended upwards with later administration. This trend could reflect a transient inflammatory response following AAV9 delivery that diminishes over time, as later-treated animals had a shorter interval between administration and tissue collection. However, differences in disease stage at treatment may also contribute.

An important future consideration for dose optimisation lies in the cellular distribution of NHE6 following AAV administration, which was not assessed in this study. Restoration of normal NHE6 levels could reflect similar expression across all cells or a wide range of expression, with some cells expressing little or no NHE6 and others expressing much higher levels. Previous studies employing single-cell and spatial transcriptomic approaches have demonstrated considerable variation in AAV-mediated transduction across CNS cell types and anatomical regions^39–41^. Consequently, restoration of total cerebellar NHE6 expression to wildtype levels does not necessarily indicate physiological expression within individual cells. Future studies integrating cell-type-specific NHE6 quantification with spatial analysis of vector distribution would help establish the proportion of cells requiring therapeutic correction, the extent of cell-to-cell variability and the upper limits of tolerable NHE6 expression. Defining these parameters will be important for identifying a dose that provides sufficient therapeutic effect while minimising the risk of excessive transgene expression.

Several limitations should be considered. First, molecular analyses were restricted to cerebellum, despite the widespread neurological phenotype of CS. Neonatal ICV may facilitate broad vector distribution within developing CNS but regional transduction efficiency is influenced by age, delivery route and vector characteristics^23^. This possibility remains speculative without comparative assessment of regional biodistribution and transgene expression in late administration of AAV9 gene replacement therapy. Second, functional efficacy was evaluated predominantly through motor phenotype, and future studies will need to consider epilepsy and intellectual phenotypes, among others. Third, AAV9 may not represent the optimal serotype capsid for *SLC9A6* replacement. Its selection provides a comparatively established and translationally relevant platform, and its expired patent reduces costs of translation. However, our previous work demonstrated sustained motor benefits following *Slc9a6* gene replacement via the engineered PhP.eB capsid at ∼5 weeks of age^15^. In contrast, while comparably timed AAV9-mediated gene replacement delayed the ataxia onset, therapeutic efficacy was fleeting, suggesting that AAV9 may not be the optimal serotype. Future studies comparing AAV9 with engineered capsids with more sustained neuronal tropism may improve therapeutic efficacy following later administration. If engineered capsids may generate a more sustained motor and/or molecular benefit, there may thus be an interplay between dose timing and the need for capsids with improved neuronal tropism. Finally, while the current study reports on 75 rats, certain comparisons are likely underpowered, particularly when multiple comparison corrections were needed. Thus, more targeted future experiments with more subjects or reduced comparisons may more conclusively resolve a number of insignificant trends in this study.

This study demonstrates that AAV9-mediated *SLC9A6* gene replacement substantially attenuates numerous motor, structural, and molecular abnormalities associated with Christianson syndrome in the *shaker* rat model, with therapeutic efficacy strongly linked to the promoter, dose, and timing of administration. Neonatal treatment produced the most pronounced and sustained benefits, including delayed onset of ataxia, preservation of cerebellar volume and improvement in disease-associated molecular markers. The lack of reduction in IBA1 and p62 upregulation despite substantial molecular rescue further highlights the importance of optimising transgene expression and evaluating treatment-associated glial and downstream lysosomal responses. Together, these findings provide further preclinical support for AAV9-mediated *SLC9A6* replacement as a potential disease-modifying therapy for CS. Future development should prioritise characterising cellular and regional transgene distribution, establishing long-term safety, and identifying approaches capable of extending treatment efficacy beyond the neonatal period, whether through engineered capsids with more extended tropism for neurons or other approaches.

## Materials and Methods

### Animal subjects and ethical approvals

All animal procedures were conducted at the Charles Perkins Centre, University of Sydney, under approval from the University of Sydney Animal Ethics Committee, protocol 2023/AE002319. A total of 75 Wistar Furth rats were included in these studies, specifically 61 *shaker* rats and 14 wildtype rats. Animals were group-housed (four per cage) under a 12:12 h light dark cycle *with ad libitum* access to food and water. For all rats enrolled in this study, *Slc9a6 g*enotyping was performed by Transnetyx (Cordova, TN, USA) using ear tissue samples collected by University of Sydney Laboratory Animal Services technical support staff. As an X-linked recessive disorder, Christianson syndrome is reported in males; however, females homozygous for the *shaker* mutation are likewise characterised full NHE6 loss of function. Thus, we utilise the term *shaker* rat to refer to both males hemizygous for the *shaker* mutation and females homozygous for the *shaker* mutation. No rats heterozygous for the *shaker* mutation were included in this study.

For the first major experimental cohort, we utilised 30 rats, consisting of 25 *shaker* rats and 5 wildtype controls. We administered AAV9 vectors via intracerebroventricular (ICV) injection to *25 shaker* rats at an average of 53 days of age, comprised of 25 *shaker* rats (7 male, 18 female), while 5 Wistar Furth wildtype males were un-injected and used as controls. *Shaker* rats were randomly assigned to five groups, with each group consisting of five rats: AAV9-SYN1-SLC9A6 at a dose of 1.0 x 10^10^ vg, AAV9-CAG-SLC9A6 at 1.0 x 10^10^ vg, AAV9-SYN1-SLC9A6 at 1.0 x 10¹¹ vg, AAV9-CAG-SLC9A6 at 1.0 × 10¹¹ vg, and AAV9-CAG-GFP at 1.0 × 10^10^ vg as surgical controls. One rat in the AAV9-CAG-SLC9A6 1.0 × 10^10^ vg group died between the 20 and 21 week motor recordings from a lung pathology identified post-mortem and deemed as unrelated to AAV administration; this group is reported with n = 4 after week 20 in the motor recordings. Notably, this animal was excluded from total ataxic burden statistical analyses.

For the second major experimental cohort, we utilised 37 rats consisting of 32 *shaker* rats and 5 wildtype controls. We aimed for 5 rats in all groups: untreated *shaker* rats, *shaker* rats treated at each of 4 days of age, 5 weeks of age, 7 weeks of age, 10 weeks of age, and 12 weeks of age, as well as wildtype controls, with all treated animals administered AAV9-SYN1-SLC9A6 at a dose of 1.0 x 10^11^ vg at their respective timepoints. While efforts were made to habituate the mother of the pups injected at postnatal day 4 to investigators, a modest rate of maternal rejection was anticipated following handling associated with AAV administration. Thus, we administered AAVs to 8 rats for the postnatal day 4 group. As only 1 pup was ultimately rejected, occurring shortly after the procedure, 7 rats were included in this group through the longitudinal study. In the second major experimental cohort, all 37 rats survived the full duration of the longitudinal study.

Finally, 8 additional age-matched rats—4 wildtype and 4 untreated *shaker*—were utilised in MRI studies in the context of the timepoint variation study.

### AAV Production

AAV vectors were designed in collaboration with the RTW Foundation, New York City, NY, USA. Human cDNA was amplified for *SLC9A6* encoding the 701 amino acid NHE6v-1 spice variant and cloned to a pAAV/SYN1-WPRE-bGHpA (utilising the neuronally targeted SYN1 promoter) or pAAV/CAG-bGHpA plasmid (utilising the ubiquitously targeted CAG promoter), with all constructs verified through sequencing. Further, eGFP was cloned into a pAAV/CAG-WPRE-bGHpA plasmid and verified through sequencing. Importantly, due to packaging size limitations, the CAG plasmid that *SLC9A6* was cloned to did not contain a woodchuck hepatitis virus posttranscriptional regulatory element, and thus CAG vs SYN1 comparisons are made within this context and may not be applicable to other disorders. Recombinant AAV particles were generated by Signanen with a titre of 2.14E13 vg/mL (SYN1), 1.10E13 vg/mL (CAG-SLCA9A6), or 3.18E13 vg/mL (CAG-eGFP). AAVs were aliquoted into 25 µL and thawed shortly prior to administration, with freeze-thaw cycles limited to 2.

### AAV Administration

For all AAV administrations other than those performed on neonatal pups, administrations were performed as previously described^15,42^. Briefly, rats were anesthetised and maintained on ∼2% isoflurane in 1 L/min oxygen. Analgesia consisted of lignocaine (2 mg/kg, local) applied at the incision site, as well as meloxicam (5 mg/kg, SC) administered daily for 3 days after surgery. Following a midline scalp incision, the skull was cleaned and dried, and craniotomies were performed at the target coordinates. Volumes injected into all cohorts were normalized to 9.09 µL of rAAV. Thus, 9.09 μL of AAV were infused at 1 μL/min via a stereotactically mounted pump, followed by a 3-min dwell time before needle withdrawal at 1.0 mm/min. Animals were given a minimum 3-day recovery period prior to motor testing.

Neonatal administration was performed on 4-day-old pups, placed on a laboratory task wipe on an aluminium tray on top of ice until unconscious. The injection site was verified and marked with surgical pen before injecting directly through the skin and skull. Following the injection the needle was left in place for ∼2 min before withdrawal. The duration of administration was kept to a minimum, and the pups were transferred to a heating pad for recovery before being returned to their home cage with the mother until the point of weaning at ∼21 days of age.

### Motor Analyses

Motor function was assessed using a custom Force Plate Actometer (FPA), modified from an original design^43^, as previously described^12,13,19,21,22^. Animals were recorded under awake, freely moving conditions weekly from 8 to 25 weeks of age for 20 min per session. Centre of mass was continuously recorded at 1000 Hz and imported into MATLAB for analysis. Straightness of gait was quantified as a proxy for gait ataxia from automatically identified rapid movements as the ratio of distance travelled to displacement as previously described^19^.

### Tissue Collection

Rats were euthanised at either 37-38 weeks of age (dose and timepoint varying study) or 29-30 weeks of age (timepoint varying study) through standard intracardiac perfusion, and brain and other tissues were collected. One half of each brain was allocated for molecular analysis, while the other for neuroimaging and histology, randomised across left vs right. In the rare instance of damage to the brain during tissue extraction, the less-damaged half was utilised for neuroimaging and histology. Tissues used for molecular analysis were flash frozen on dry ice and homogenized prior to protein extraction using RIPA Buffer with protease and phosphatase inhibitors. The contralateral hemisphere was post-fixed in 3% paraformaldehyde (PFA) for 48 h followed by cryoprotection and long-term storage in PBS with 0.05% sodium azide at 4°C. Other organs were collected and flash frozen for future study, including brain stem, lungs, heart, liver, spleen, kidney, and hind limbs.

### Protein Analysis and Antibodies

Western blots were performed similarly to previously published protocols ^42,44^, with protein quantification performed relative to GAPDH or Beta-actin loading controls. The following primary antibodies were used: Anti-SLC9A6 [(1:1000), GeneTex Cat# GTX32878], GAPDH [(1:5000), Cell Signalling, rabbit mAb #2118], ACTB [(1:10,000), Sigma-Aldrich mouse mAb #A3854], CALB1 [(1:5000), Sigma-Aldrich, mouse mAb #C9848], SQSTM1/p62 [(1:1000), Sigma-Aldrich rabbit pAb #P0068]. RGS8 [(1:1000), Novus Biologicals, #NBP2-20153)], PCP2 [(1:1000), Santa Cruz Biotechnology, rabbit mAB # SC-137064], GFAP [(1:2000), ThermoFisher Scientific, rat mAb #130300], IBA-1 [(1:1000), Abcam Australia, rabbit mAb # AB178846]. Secondary antibodies used included HRP-Goat Anti-Rabbit IgG (H+L) [(1:5000), Jackson ImmunoResearch Laboratories, Cat# 111-035-144] and HRP-Horse Anti-Mouse IgG (H+L) [(1:5000), Vector Laboratories, Cat# PI-2000].

Notably, in the instance of NHE6 blotting in the timepoint variation experiment, it was not possible to resolve both rats with maximal treatment effect (increased from wildtype) and minimal treatment effect (on the order of a few percent of wildtype) on the same exposure without oversaturation of the strongest bands. Thus, blots were imaged with both short and long exposures and P4-treated rats were quantified and normalised to wildtype on short exposure blots, while remaining groups were quantified and normalised to wildtype on long exposure blots.

### Magnetic Resonance Imaging (MRI)

MRI acquisition was performed at the Sydney Imaging Facility, University of Sydney, using a T-weighted 3D sequence to obtain high-resolution anatomical images of the whole brain and cerebellum. Scans were acquired with a field of view (FOV) of 12 × 10 × 16–24 mm³ and a matrix size of 128 × 100 × 160–240, yielding an isotropic voxel size of 100 µm³. Each scan was averaged over 8–16 acquisitions to optimize the signal-to-noise ratio.

Volumetric analysis of the cerebellum was performed using 3D Slicer (version 5.10.0). A master segmentation was registered for both left and right hemispheres, and a Python-based registration pipeline was made to morph master segmentation on the input scan. This was followed by investigator-blinded manual segmentation for anatomical precision and accurate capture of the cerebellum across axial, coronal and sagittal planes. Following blinded manual refinement, hemi-cerebellar volume was quantified using segment statistics in 3D Slicer and reported in mm^3^.

Seven brains were excluded from MRI imaging in the dose timepoint analysis experiment. First, one P4-treated rat was deemed to have had its cerebellum damaged in processing prior to imaging. Second, 2 untreated *shaker* rats and 4 of the 5 W12-treated rats were deemed to have been poorly post-fixed and unsuitable for imaging. In the absence of additional W12-treated rats and only one subject imaged, this group was excluded from analyses, although we note that the hemicerebellar volume of this animal imaged was consistent with those of other late-stage treatment rats. Notably, given the availability and parallel processing of 8 additional brains taken from age-matched rats—4 wildtype and 4 untreated *shaker*—at the point of euthanasia of timepoint varying experimental rats, these brains were included to increase statistical power of the study.

### Statistical analysis

Statistical analyses were performed via several approaches. First, longitudinal motor outcomes were analysed using linear mixed-effects models to assess the effects of treatment group, age and their interaction, accounting for repeated measurements at a given timepoint within individual animals; in these analyses, statistics are reported through Joint Wald F tests. Kaplan-Meier curves for motor thresholds were evaluated using χ^2^ testing for both group-wise and post-hoc differences, comparing each other group to untreated *shaker* rats, with a Holm-Bonferroni correction applied in the context of post-hoc testing. Finally, total ataxic burden was quantified through simplification of the longitudinal data to a single average severity score, with statistical analyses performed via one-way ANOVA with Holm-Bonferroni post-hoc corrections, with all other groups compared to untreated *shaker* rats in both studies and P4-treated rats compared to wildtype rats in the second experiment. Notably, in the total ataxic burden analyses, one rat was excluded from this analysis in the CAG 1 x 10^10^ treatment due to early fatality deemed unrelated to treatment.

For each western blotting experiment, a one-tailed heteroscedastic (Welch’s) t-test was first performed to determine whether a significant difference was present between wildtype rats and untreated *shaker* rats. If a significant difference was not found between wildtype rats and *shaker* rats, further statistical testing was not performed. If a significant difference was found, a one-way ANOVA test was then performed to determine whether group-based differences were found. If the ANOVA test yielded statistical significance, then post-hoc tests were performed for individual comparisons beyond wildtype rats versus untreated *shaker* rats. When more specific hypotheses were informed by prior work (specifically, NHE6, PCP2, CALB1, and RGS8), Holm-Bonferroni-corrected one-tailed heteroscedastic (Welch’s) post-hoc t-tests were performed: wildtype rats versus *shaker* rats treated at P4, and all treatment groups (P4, week 5, week 7, week 10, and week 12) individually versus untreated *shaker* rats. When we were evaluating specific proteins for the first time, we instead performed Tukey’s Honestly Significant Difference testing to compare each group against all other groups.

For structural MRI experiments, first, a one-tailed heteroscedastic (Welch’s) t-test was performed to determine whether a significant difference was present between wildtype rats and untreated *shaker* rats. In both experiments, given that a significant difference was found, a one-way ANOVA test was then performed to determine whether group-based differences were present. Again, if a significant difference was found in both cases, Holm-Bonferroni-corrected hesteroscedastic (Welch’s) t-tests were performed as post-hoc testing to evaluate each treated group separately versus untreated *shaker* rats from the corresponding experiment, with one further Holm Bonferroni-corrected one-tailed heteroscedastic (Welch’s) t-test to compare P4-treated *shaker* rats with corresponding wildtype rats.

We utilised cohorts with temporally different AAV dosing (P4, W5, W7, W10, W12) to calculate decay and accumulation properties of molecular markers and other metrics, with sex as a covariate and Holm-Bonferroni was applied as a post-hoc correction in every test. First, we z-scored metrics and fit linear slopes against dosing age. Regression for linear and exponential trajectories used standard forms, and we reported R^2^ values to quantify goodness of each fit and extracted half-life values from exponential fits. We used fold change relative to the WT mean to quantify how inflammation markers increased as a multiplicative factor over wildtype levels and fit a regression on a log2 scale to quantify weekly change.

Heatmap correlations were calculated using Pearson’s r analyses when standard assumptions were met regarding statistical distributions and residuals; when standard assumptions were not met, most typically through significant non-normality of distributions, Spearman’s ρ was calculated instead. We used Holm-Bonferroni to correct for multiple comparisons within each set of planned tests but used Benjamini-Hochberg correction for exploratory marker-motor correlations across weeks. We ran a principal component analysis (PCA) on standardised molecular markers to identify patterns of shared variance among them. We used a multivariate, linear model to predict final ataxia burden using all molecular markers and sex, and we reported raw and adjusted R^2^ to quantify goodness of fit. We reported all coefficients from linear models with 95% confidence intervals. We used relative importance analysis to rank variables in order of most to least explained variables, shown with 95% confidence intervals, and used the rankings to select three markers a simplified linear model with sex as a covariate. A nested F-test compared a full model of ataxic burden containing all markers and with a sex-only baseline model to evaluate the time point when cohorts diverse in weekly ataxic burden.

Aside from any deviations described above, bar plots show means, while error clouds and error bars represent standard error of the mean. In all experiments, statistical significance was defined as p< 0.05 following post-hoc testing if applicable, with significance shown on figures as * for p<.05, ** for p<.01, and *** for p<.001, once again following post-hoc testing if applicable. P-values are reportedly exactly for values above p≥.001, whilst values below p<.001 are reported as such.

## Notes

### Competing Interest Statement

The authors have declared no competing interest.

## References

1. Christianson AL, Stevenson RE, Van der Meyden CH, et al. X linked severe mental retardation, craniofacial dysmorphology, epilepsy, ophthalmoplegia, and cerebellar atrophy in a large South African kindred is localised to Xq24-q27. J Med Genet. 1999;36(10):759–766. doi:10.1136/jmg.36.10.759

2. Gilfillan GD, Selmer KK, Roxrud I, et al. SLC9A6 Mutations Cause X-Linked Mental Retardation, Microcephaly, Epilepsy, and Ataxia, a Phenotype Mimicking Angelman Syndrome. The American Journal of Human Genetics. 2008;82(4):1003–1010. 10.1016/j.ajhg.2008.01.013

3. Pescosolido MF, Ouyang Q, Liu JS, Morrow EM. Loss of Christianson syndrome Na+/H+ exchanger 6 (NHE6) causes abnormal endosome maturation and trafficking underlying lysosome dysfunction in neurons. Journal of Neuroscience. 2021;41(44):9235–9256. doi:10.1523/JNEUROSCI.1244-20.2021

4. Ouyang Q, Lizarraga SB, Schmidt M, et al. Christianson Syndrome Protein NHE6 Modulates TrkB Endosomal Signaling Required for Neuronal Circuit Development. Neuron. 2013;80(1):97–112. 10.1016/j.neuron.2013.07.043

5. Kavanaugh BC, Elacio J, Best CR, et al. Christianson syndrome across the lifespan: genetic mutations and longitudinal study in children, adolescents, and adults. J Med Genet. 2024;61(11):1031–1039. doi:10.1136/jmg-2024-109973

6. Premachandran S, Ocay DD, Beaulieu C, et al. Pain experience of children with Christianson syndrome. Pain. 2025;166(7):1610–1621. doi:10.1097/j.pain.0000000000003522

7. Regina L, Yates-Siilata MC, Woods K, Tolbert L. Preliminary characterization of hereditary cerebellar ataxia in rats. Lab Anim Sci. 1992;42(1):19–26.

8. Clark BR, LaRegina M, Tolbert DL. X-linked transmission of the shaker mutation in rats with hereditary Purkinje cell degeneration and ataxia. Brain Res. 2000;858(2):264–273. doi:10.1016/s0006-8993(99)02415-4

9. Wolf LW, LaRegina MC, Tolbert DL. A behavioral study of the development of hereditary cerebellar ataxia in the shaker rat mutant. Behavioural Brain Research. 1996;75(1):67–81. 10.1016/0166-4328(96)00159-3

10. Tolbert DL, Ewald M, Gutting J, La Regina MC. Spatial and temporal pattern of purkinje cell degeneration in shaker mutant rats with hereditary cerebellar ataxia. Journal of Comparative Neurology. 1995;355(4):490–507. 10.1002/cne.903550403

11. Tolbert DL, Clark BR. GDNF and IGF-I trophic factors delay hereditary Purkinje cell degeneration and the progression of gait ataxia. Exp Neurol. 2003;183(1):205–219. 10.1016/S0014-4886(03)00172-9

12. 42nd Annual International Conferences of the IEEE Engineering in Medicine and Biology Society : “Enabling Innovative Technologies for Global Healthcare” : 20-24 July 2020, Montreal, Canada. IEEE; 2020.

13. Figueroa Kp, Anderson Cj, Paul S, Et Al. Slc9A6 Mutation Causes Purkinje Cell Loss And Ataxia In The Shaker Rat. Hum Mol Genet. 2023;32(10):1647–1659. doi:10.1093/hmg/ddad004

14. Figueroa KP, Paul S, Calì T, et al. Spontaneous shaker rat mutant – a new model for X-linked tremor-ataxia. Dis Model Mech. Published online 2016. doi:10.1242/dmm.022848

15. Anderson CJ, Figueroa KP, Paul S, et al. Viral Vector-Mediated Slc9A6 Gene Replacement Reduces Cerebellar Motor And Molecular Abnormalities In The Shaker Rat Model Of Christianson Syndrome. Hum Mol Genet. 2026;35(6). doi:10.1093/hmg/ddag021

16. Ling Q, Herstine JA, Bradbury A, Gray SJ. AAV-based in vivo gene therapy for neurological disorders. Nat Rev Drug Discov. Nature Research. 2023;22(10):789–806. doi:10.1038/s41573-023-00766-7

17. Byrne BJ, Flanigan KM, Matesanz SE, et al. Current clinical applications of AAV-mediated gene therapy. Molecular Therapy. Cell Press. 2025;33(6):2479–2516. doi:10.1016/j.ymthe.2025.04.045

18. Anderson CJ, Cadeddu R, Anderson DN, et al. A novel naïve Bayes approach to identifying grooming behaviors in the force-plate actometric platform. J Neurosci Methods. 2024;403. doi:10.1016/j.jneumeth.2023.110026

19. Anderson CJ, Figueroa KP, Dorval AD, Pulst SM. Deep cerebellar stimulation reduces ataxic motor symptoms in the shaker rat. Ann Neurol. 2019;85(5):681–690. 10.1002/ana.25464

20. Paul S, Dansithong W, Figueroa KP, Scoles DR, Pulst SM. Staufen1 links RNA stress granules and autophagy in a model of neurodegeneration. Nature Communications. 2018;9(1). doi:10.1038/s41467-018-06041-3

21. Gray SJ. Timing of Gene Therapy Interventions: The Earlier, the Better. Molecular Therapy. Nature Publishing Group. 2016;24(6):1017–1018. doi:10.1038/mt.2016.20

22. McLean JR, Smith GA, Rocha EM, et al. Widespread neuron-specific transgene expression in brain and spinal cord following synapsin promoter-driven AAV9 neonatal intracerebroventricular injection. Neurosci Lett. 2014;576:73–78. doi:10.1016/j.neulet.2014.05.044

23. Gholizadeh S, Tharmalingam S, MacAldaz ME, Hampson DR. Transduction of the central nervous system after intracerebroventricular injection of adeno-associated viral vectors in neonatal and juvenile mice. Hum Gene Ther Methods. 2013;24(4):205–213. doi:10.1089/hgtb.2013.076

24. Galvan A, Petkau TL, Hill AM, et al. Intracerebroventricular Administration of AAV9-PHP.B SYN1-EmGFP Induces Widespread Transgene Expression in the Mouse and Monkey Central Nervous System. Hum Gene Ther. 2021;32(11-12):599–615. doi:10.1089/hum.2020.301

25. Hughes MP, Smith DA, Morris L, et al. AAV9 intracerebroventricular gene therapy improves lifespan, locomotor function and pathology in a mouse model of Niemann-Pick type C1 disease. Hum Mol Genet. 2018;27(17):3079–3098. doi:10.1093/hmg/ddy212

26. Hatten ME, Alder’ J, Zimmerman’ K, He&z3 N. Genes Involved in Cerebellar Cell Differentiation Specification And.

27. Altman J. Postnatal development of the cerebellar cortex in the rat. I. The external germinal layer and the transitional molecular layer. Journal of Comparative Neurology. 1972;145(3):353–397. doi:10.1002/cne.901450305

28. Foust KD, Nurre E, Montgomery CL, Hernandez A, Chan CM, Kaspar BK. Intravascular AAV9 preferentially targets neonatal neurons and adult astrocytes. Nat Biotechnol. 2009;27(1):59–65. doi:10.1038/nbt.1515

29. Sługocka A, Wiaderkiewicz J, Barski JJ. Genetic Targeting in Cerebellar Purkinje Cells: an Update. Cerebellum. Springer New York LLC. 2017;16(1):191–202. doi:10.1007/s12311-016-0770-4

30. Zhang X, Zhang H, Oberdick J. C Onservation of the Developmentally Regulated Dendritic Localization of a Purkinje Cell-Specific MRNA That Encodes a G-Protein Modulator: Comparison of Rodent and Human Pcp2(L7) Gene Structure and Expression. Vol 105. 2002. www.elsevier.com/locate/bres

31. Garcia-Segura LM, Baetens D, Roth J, Norman AW, Orci L. Immunohistochemical Mapping of Calcium-Binding Protein Immunoreactivity in the Rat Central Nervous System. Vol 296. 1984.

32. Arnold DB, Heintz N. A Calcium Responsive Element That Regulates Expression of Two Calcium Binding Proteins in Purkinje Cells (Transcriptionbiolistic TransfectionCa 2 Responsescalbindincalmodulin II). Vol 94. 1997. www.pnas.org.

33. Barski JJ, Hartmann J, Rose CR, et al. Calbindin in Cerebellar Purkinje Cells Is a Critical Determinant of the Precision of Motor Coordination. 2003.

34. Xu M, Ouyang Q, Gong J, et al. Mixed Neurodevelopmental and Neurodegenerative Pathology in Nhe6-Null Mouse Model of Christianson Syndrome. eNeuro. 2017;4(6):ENEURO.0388-17.2017. doi:10.1523/ENEURO.0388-17.2017

35. Lee Y, Miller MR, Fernandez MA, et al. Early lysosome defects precede neurodegeneration with amyloid-β and tau aggregation in NHE6-null rat brain. Brain. 2022;145(9):3187–3202. doi:10.1093/brain/awab467

36. Fernandez MA, Bah F, Ma L, et al. Loss of endosomal exchanger NHE6 leads to pathological changes in tau in human neurons. Stem Cell Reports. 2022;17(9):2111–2126. doi:10.1016/j.stemcr.2022.08.001

37. Samaranch L, Sebastian WS, Kells AP, et al. AAV9-mediated expression of a non-self protein in nonhuman primate central nervous system triggers widespread neuroinflammation driven by antigen-presenting cell transduction. Molecular Therapy. 2014;22(2):329–337. doi:10.1038/mt.2013.266

38. Ciesielska A, Hadaczek P, Mittermeyer G, et al. Cerebral infusion of AAV9 vector-encoding non-self proteins can elicit cell-mediated immune responses. Molecular Therapy. 2013;21(1):158–166. doi:10.1038/mt.2012.167

39. Brown D, Altermatt M, Dobreva T, et al. Deep Parallel Characterization of AAV Tropism and AAV-Mediated Transcriptional Changes via Single-Cell RNA Sequencing. Front Immunol. 2021;12. doi:10.3389/fimmu.2021.730825

40. Jang MJ, Coughlin GM, Jackson CR, et al. Spatial transcriptomics for profiling the tropism of viral vectors in tissues. Nat Biotechnol. 2023;41(9):1272–1286. doi:10.1038/s41587-022-01648-w

41. Fouani Y, Oti M, Jarosch S, et al. Spatial mapping of barcoded, brain-tropic AAVs using multiplexed RNA in situ hybridization. Molecular Therapy Advances. 2026;34(1). doi:10.1016/j.omta.2026.201693

42. Figueroa KP, Anderson CJ, Paul S, et al. Slc9a6 mutation causes Purkinje cell loss and ataxia in the shaker rat. Hum Mol Genet. 2023;32(10):1647–1659. doi:10.1093/hmg/ddad004

43. Fowler SC, Birkestrand BR, Chen R, et al. A Force-Plate Actometer for Quantitating Rodent Behaviors: Illustrative Data on Locomotion, Rotation, Spatial Patterning, Stereotypies, and Tremor. Vol 107. 2001. www.elsevier.com/locate/jneumeth

44. Paul S, Dansithong W, Figueroa KP, Scoles DR, Pulst SM. Staufen1 links RNA stress granules and autophagy in a model of neurodegeneration. Nat Commun. 2018;9(1):3648. doi:10.1038/s41467-018-06041-3

